# Soil viruses are underexplored players in ecosystem carbon processing

**DOI:** 10.1101/338103

**Authors:** Gareth Trubl, Ho Bin Jang, Simon Roux, Joanne B. Emerson, Natalie Solonenko, Dean R. Vik, Lindsey Solden, Jared Ellenbogen, Alexander T. Runyon, Benjamin Bolduc, Ben J. Woodcroft, Scott R. Saleska, Gene W. Tyson, Kelly C. Wrighton, Matthew B. Sullivan, Virginia I. Rich

## Abstract

Rapidly thawing permafrost harbors ~30–50% of global soil carbon, and the fate of this carbon remains unknown. Microorganisms will play a central role in its fate, and their viruses could modulate that impact via induced mortality and metabolic controls. Because of the challenges of recovering viruses from soils, little is known about soil viruses or their role(s) in microbial biogeochemical cycling. Here, we describe 53 viral populations (vOTUs) recovered from seven quantitatively-derived (i.e. not multiple-displacement-amplified) viral-particle metagenomes (viromes) along a permafrost thaw gradient. Only 15% of these vOTUs had genetic similarity to publicly available viruses in the RefSeq database, and ~30% of the genes could be annotated, supporting the concept of soils as reservoirs of substantial undescribed viral genetic diversity. The vOTUs exhibited distinct ecology, with dramatically different distributions along the thaw gradient habitats, and a shift from soil-virus-like assemblages in the dry palsas to aquatic-virus-like in the inundated fen. Seventeen vOTUs were linked to microbial hosts (*in silico*), implicating viruses in infecting abundant microbial lineages from *Acidobacteria, Verrucomicrobia*, and *Deltaproteoacteria*, including those encoding key biogeochemical functions such as organic matter degradation. Thirty-one auxiliary metabolic genes (AMGs) were identified, and suggested viral-mediated modulation of central carbon metabolism, soil organic matter degradation, polysaccharide-binding, and regulation of sporulation. Together these findings suggest that these soil viruses have distinct ecology, impact host-mediated biogeochemistry, and likely impact ecosystem function in the rapidly changing Arctic.

## Importance

This work is part of a 10-year project to examine thawing permafrost peatlands, and is the first virome-particle-based approach to characterize viruses in these systems. This method yielded >2-fold more viral populations (vOTUs) per gigabase of metagenome than vOTUs derived from bulk-soil metagenomes from the same site (Emerson et al. *in press*, Nature Microbiology). We compared the ecology of the recovered vOTUs along a permafrost thaw gradient, and found: (1) habitat specificity, (2) a shift in viral community identity from soil-like to aquatic-like viruses, (3) infection of dominant microbial hosts, and (4) encoding of host metabolic genes. These vOTUs can impact ecosystem carbon processing via top-down (inferred from lysing dominant microbial hosts) and bottom-up (inferred from encoding auxiliary metabolic genes) controls. This work serves as a foundation upon which future studies can build upon to increase our understanding of the soil virosphere and how viruses affect soil ecosystem services.

## Introduction

Anthropogenic climate change is elevating global temperatures, most rapidly at the poles (1). High-latitude perennially-frozen ground, i.e. permafrost, stores 30–50% of global soil carbon (C; ~1300 Pg; 2, 3) and is thawing at a rate of ≥1 cm of depth yr^−1^ (4, 5). Climate feedbacks from permafrost habitats are poorly constrained in global climate change models (1, 6), due to the uncertainty of the magnitude and nature of carbon dioxide (CO_2_) or methane (CH_4_) release. A model ecosystem for studying the impacts of thaw in a high-C peatland setting is Stordalen Mire, in Arctic Sweden, which is at the southern edge of current permafrost extent (7). The Mire contains a mosaic of thaw stages (8), from intact permafrost palsas, to partially-thawed moss-dominated bogs, to fully-thawed sedge-dominated fens (9–12). Thaw shifts hydrology (13), altering plant communities (12), and shifting belowground organic matter (OM) towards more labile forms (10, 12), with concomitant shifts in microbiota (14–16), and C gas release (7, 9, 17–19). Of particular note is the thaw-associated increase in CH_4_ emissions, due to its 33-times greater climate forcing potential than CO_2_ (per kg, at a 100-year time-scale; 20), and the associated shifts in key methanogens. These include novel methanogenic lineages (14) with high predictive value for the character of the emitted CH_4_ (11). More finely resolving the drivers of C cycling, including microbiota, in these dynamically changing habitats can increase model accuracy (21) to allow a better prediction of greenhouse gas emissions in the future.

Given the central role of microbes to C processing in these systems, it is likely that viruses infecting these microbes impact C cycling, as has been robustly observed in marine systems (22–27). Marine viruses lyse ~one-third of ocean microorganisms day^−1^, liberating C and nutrients at the global scale (22–24, 28), and viruses have been identified as one of the top predictors of C flux to the deep ocean (29). Viruses can also impact C cycling by metabolically reprogramming their hosts, via the expression of viral-encoded “auxiliary metabolic genes” (AMGs; 28, 30) including those involved in marine C processing (31–35). In contrast, very little is known about soil virus roles in C processing, or indeed about soil viruses generally. Soils’ heterogeneity in texture, mineral composition, and OM content results in significant inconsistency of yields from standard virus ‘capture’ methods (36–39). While many soils contain large numbers of viral particles (10^7^–10^9^ virus particles per gram of soil; 37, 40–42), knowledge of soil viral ecology has come mainly from the fraction that desorb easily from soils (<10% in 43) and the much smaller subset that have been isolated (44).

One approach to studying soil viruses has been to bypasses the separation of viral particles, by identifying viruses from bulk-soil metagenomes; these are commonly referred to as microbial metagenomes but contain sequences of diverse origin, including proviruses and infecting viruses. Using this approach, several recent studies have powerfully expanded our knowledge of soil viruses and have highlighted the magnitude of genetic novelty these entities may represent. An analysis of 3,042 publicly-available assembled metagenomes spanning 10 ecotypes (19% from soils) increased by 16-fold the total number of known viral genes, doubled the number of microbial phyla with evidence of viral infection, and revealed that the vast majority of viruses appeared to be habitat-specific (45). This approach was also applied to 178 metagenomes from the thawing permafrost gradient of Stordalen Mire (46), where viral linkages to potential hosts were appreciably advanced by the parallel recovery of 1,529 microbial metagenome-assembled genomes (MAGs; 16). This effort recovered ~2000 thaw-gradient viruses, more than doubling known viral genera in Refseq, identified linkages to abundant microbial hosts encoding important C-processing metabolisms such as methanogenesis, and demonstrated that CH4 dynamics was best predicted by viruses of methanogens and methanotrophs (46). Viral analyses of bulk-soil metagenomes have thus powerfully expanded knowledge of soil viruses and highlighted the large amount of genetic novelty they represent. However this approach is by nature inefficient at capturing viral signal, with typically <2% of reads identified as viral (46, 47). The small amount of viral DNA present in bulk-soil extracts can lead to poor or no assembly of viral sequences in the resulting metagenomes, and omission from downstream analyses (discussed further in 37, 39, 48, 49). In addition, viruses that are captured in bulk-soil metagenomes likely represent a subset of the viral community, since >90% of free viruses adsorb to soil (43), and so depending on the specific soil, communities, and extraction conditions, bulk-soil metagenomes are likely be depleted for some free viruses and enriched for actively reproducing and temperate viruses.

Examination of free viruses, while potentially a more efficient and comprehensive approach to soil viral ecology, requires optimized methods to resuspend them (50). Researchers have pursued optimized viral resuspension methods for specific soil types, and metagenomically sequenced the recovered viral particles, generating *viromes.* In marine systems, viral ecology has relied heavily on viromes, since the leading viral particle capture method is broadly applicable, highly efficient, and relatively inexpensive (51), with now relatively well-established downstream pipelines for quantitative sample-to-sequence (52) and sequence-to-ecological-inference (53, 54) processing, collectively resulting in great advances in marine viromics (55). Due to the requirement of habitat-specific resuspension optimization, soil viromics is in its early stages. In addition, because particle yields are typically low, most soil virome studies have amplified extracted viral DNA using multiple displacement amplification, which renders the datasets both stochastically and systematically biased and non-quantitative (53, 56–61). The few polar soil viromes have been from Antarctic soils, and further demonstrated the genetic novelty of this gene pool, while suggesting resident viral communities were dominated by tailed-viruses, had high habitat specificity, and were structured by pH (62–64).

Having previously optimized viral resuspension methods for the active layer of the permafrost thaw gradient in the Stordalen Mire (41), here we sequenced and analyzed a portion of the viruses recovered from that optimization effort, with no amplification beyond that minor, quantitative form inherent to sequencing library preparation. The seven resulting viromes yielded 378 genuine viral contigs, 53 of which could be classified as vOTUs (approximately representing species-level taxonomy; 65). The goal of this effort was to efficiently target viral particle genomes via viromes from Stordalen Mire, investigate their ecology and potential impacts on C processing using a variety of approaches, and compare the findings to that of viral analyses of bulk-soil-metagenomes from Emerson et al. (46).

## Results and Discussion

### Viruses in complex soils

Using recently develop bioinformatics tools to characterize viruses from three different habitats along a permafrost thaw gradient, viral particles were purified from active layer soil samples (i.e. samples from the upper, unfrozen portion of the soil column) via a previously-optimized method tailored for these soils (41; Fig. 1). DNA from viral particles was extracted and sequenced, to produce seven Stordalen Mire viromes (Table 1), spanning palsas (underlain by intact permafrost), bogs (here underlain by partially-thawed permafrost), and fens (where permafrost has thawed entirely). The viromes ranged in size from 2-26 million reads, with an average of 18% of the reads assembling into 28,025 total contigs across the dataset. Among these, VirSorter predicted that 393 contigs were viruses (VirSorter categories 1, 2, and 3; per 66; see Methods; Table 2). After manual inspection, three putative plasmids were identified and removed (i.e. contigs 5, 394, and 3167; Table 2), along with two putatively archaeal viruses (vOTUs 165 and 225; analyzed separately, see supplementary information). Finally, ten additional contigs that did not meet our threshold for read recruitment (i.e. 90% average nucleotide identity across 75% of contig covered) were removed, resulting in 378 putative virus sequences (Table 2). Of these, 53 bacteriophage (phage) were considered well-sampled ‘viral populations’ (54) also known as viral operational taxonomic units (vOTUs) as they had contig lengths ≥10 kb (average 19.6 kb, range: 10.3 kb–129.6 kb), were most robustly viral (VirSorter category 1 or 2; 66), and were relatively well-covered contigs (averaged 74x coverage, Table 1). These 53 viral populations are the basis for the analyses in this paper due to their genome sizes, which allowed for more reliable taxonomic, functional, and host assignments, and fragment recruitment.

**Figure 1.**
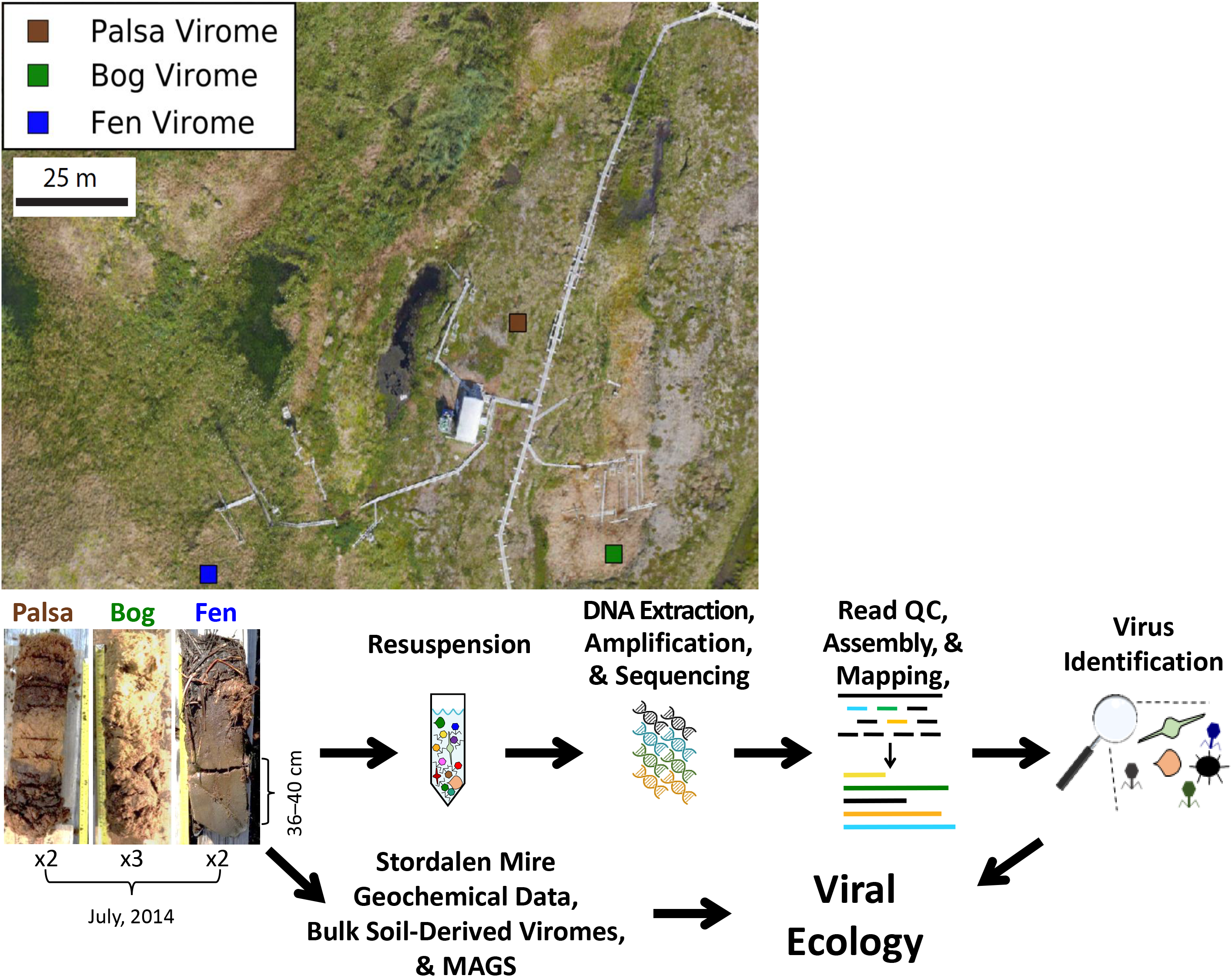
Overview of sample-to-ecology methods pipeline. Sampling of the thaw chronosequence at Stordalen Mire (68°21 N, 19°03 E, 359 m a.s.l.). The underlying image was collected via unmanned aerial vehicle (UAV) and extensively manually curated for GPS accuracy (generated by Dr. Michael Palace). Sampling locations were mapped onto this image based on their GPS coordinates. Soil cores were taken in July of 2014. Viruses were resuspended as previously described in Trubl et al. (41). Viromes were generated using samples from 36–40 cm. Identified vOTUs were further characterized using geochemical data and metagenome-assembled genomes (MAGs; 16) from Stordalen Mire. Additionally, these vOTUs were compared to the vOTUs from bulk-soil-derived viromes (46).

There is no universal marker gene (analogous to the 16S rRNA gene in microbes) to provide taxonomic information for viruses. We therefore applied a gene-sharing network where nodes were genomes and edges between nodes indicated the gene content similarities, and accommodating fragmented genomes of varying sizes (67–72). In such networks, viruses sharing a high number of genes localize into viral clusters (VCs) which represent approximately genus-level taxonomy (69, 72). We represented relationships across the 53 vOTUs with 2,010 known bacterial and archaeal viruses (RefSeq, version 75) as a weighted network (Fig. 2). Only 15% of the Mire vOTUs had similarity to RefSeq viruses (Fig. 2). Three vOTUs fell into 3 VCs comprised of viruses belonging to the *Fellxounavlrinae* and *Vequintavirinae* (VC10), *Tevenvirinae* and *Eucampyvirinae* (VC3), and the *Bcep22virus*, *F116virus* and *Kpp25virus* (VC4) (Fig. 2). Corroborating its taxonomic assignment by clustering, vOTU_4 contained two marker genes (i.e., major capsid protein and baseplate protein) specific for the *Felixounavirinae* and *Vequintavirinae* viruses (73), phylogenetic analysis of which indicated a close relationship of vOTU_4 to the *Cr3virus* within the *Vequintavirinae* (Fig. S2). The other five populations that clustered with RefSeq viruses were each found in different clusters with taxonomically unclassified viruses (Fig.2). Viruses derived from the dry palsa clustered with soil-derived RefSeq viruses, while those from the bog clustered with a mixture of soil and aquatic RefSeq viruses, and those from the fen clustered mainly with aquatic viruses (Fig. 2). Though of limited power due to small numbers, this suggests some conservation of habitat preference within genotypic clusters, which has also been observed in marine viruses with only ~4% of VCs being globally ubiquitous (70). Most (~85%) of the Mire vOTUs were unlinked to RefSeq viruses, with 41 vOTUs having no close relatives (i.e. singletons), and the remaining 4 vOTUs clustering in doubletons. This separation between a large fraction of the Mire vOTUs and known viruses is due to a limited number of common genes between them, i.e. ~70% of the total proteins in these viromes are unique (Fig. 2), reflecting the relative novelty of these viruses and the undersampling of soil viruses (39).

**Figure 2.**
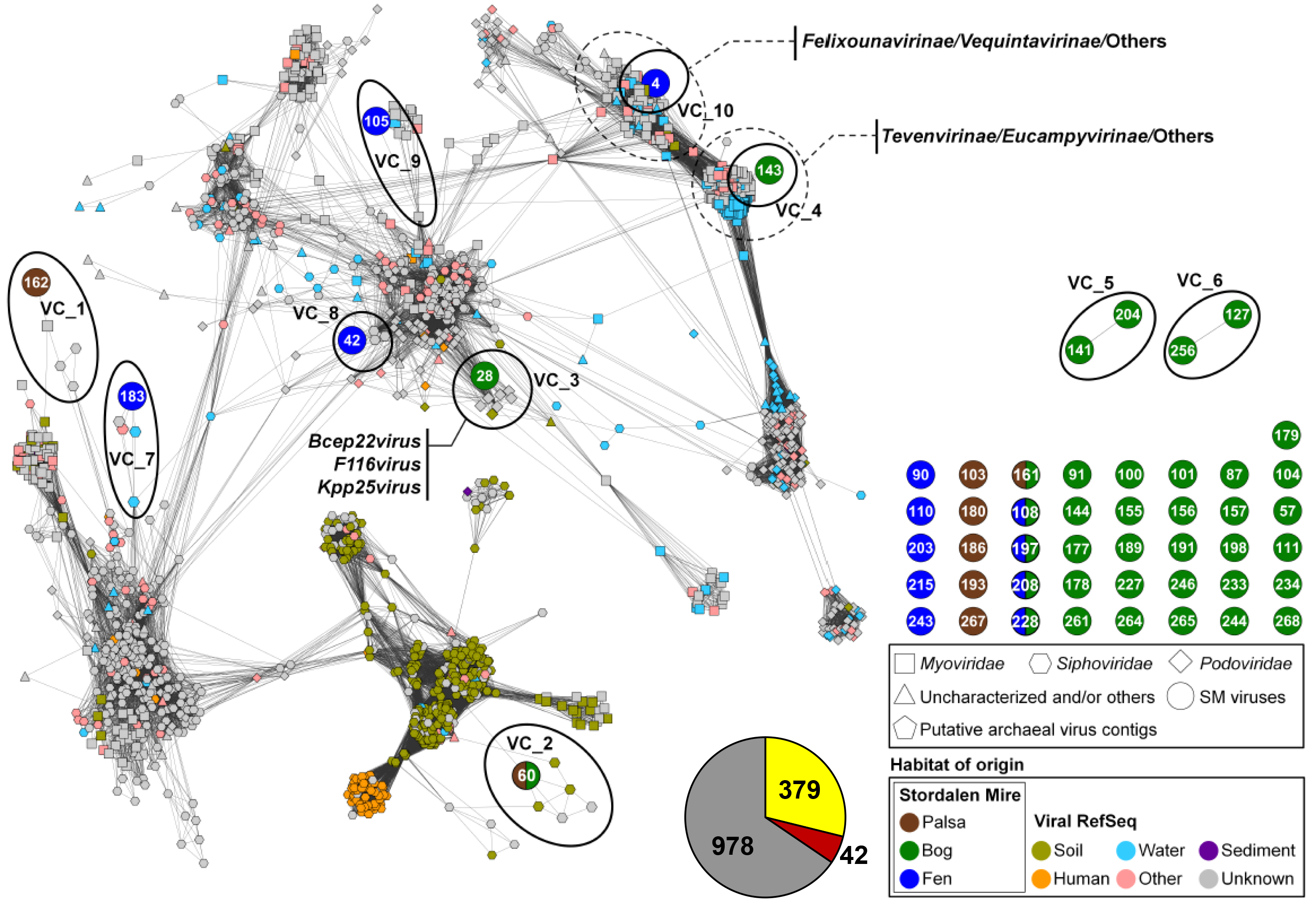
Relating Stordalen Mire viruses to known viral sequence space. Clustering of recovered vOTUs with all RefSeq (v 75) viral genomes or genome fragments with genetic connectivity to these data. Shapes indicate major viral families, and RefSeq sequences only indirectly linked to these data are in gray. The contig numbers are shown within circles. Each node is depicted as a different shape, representing viruses belonging to *Myoviridae* (rectangle), *Podoviridae* (diamond), *Siphoviridae* (hexagon), or uncharacterized viruses (triangle) and viral contigs (circle). Edges (lines) between nodes indicate statistically weighted pairwise similarity scores (see Methods) of ≥1. Color denotes habitat of origin, with “other” encompassing wastewater, sewage, feces, and plant material. Contig-encompassing viral clusters are encircled by a solid line (slightly off because it’s a 2-dimensional representation of a 3-D space). Dashed lines indicate two network regions of consistent known taxonomy, allowing assignment of contigs 4, 143, and 28. The pie chart represents the number of the Stordalen Mire viral proteins (i) that are recovered by protein clusters (PCs) (yellow and red) and singletons (gray) and (ii) that are shared with RefSeq viruses (yellow) or not (red and gray). Proteins of viral genomes/vOTUs in the dataset were grouped into PCs through all-to-all BlastP comparisons (E-value cut-off <10^−4^) followed by Markov clustering algorithm-based clustering (see Materials and Methods). Proteins that were not grouped into PCs are designated as singletons.

Annotation of the 53 vOTUs resulted in only ~30% of the genes being annotated, which is not atypical; >60% of genes encoded in uncultivated viruses have typically been classified as unknown in other studies (46, 66, 74–78). Of genes with annotations, we first considered those involved in lysogeny, to provide insight into the viruses’ replication cycle. Only three viruses encoded an integrase gene (other characteristic lysogeny genes were not detected; 79, 80; Table S1), suggesting they could be temperate viruses, two of which were from the bog habitat. It had been proposed that since soils are structured and considered harsh environments, a majority of soil viruses would be temperate viruses (81). Although our dataset is small, a dominance of temperate viruses is not observed here. We hypothesize that the low encounter rate produced by the highly structured soil environment could, rather than selecting for temperate phage, select for efficient virulent viruses (concept derived from 82–84). Recent analyses of the viral signal mined from bulk-soil metagenomes from this site provides more evidence for our hypothesis of efficient virulent viruses, because >50% of the identified viruses were likely not temperate (based on the fact they were not detected as prophage; 46). As a more comprehensive portrait of soil viruses grows, spanning various habitats, this hypothesis can be further tested. Beyond integrase genes, the remaining annotated genes spanned known viral genes and host-like genes. Viral genes included those involved in structure and replication, and their taxonomic affiliations were unknown or highly variable, supporting the quite limited affiliation of these vOTUs with known viruses. Host-like genes included AMGs, which are described in greater detail in the next section.

### Host-linked viruses are predicted to infect key C cycling microbes

In order to examine these viruses’ impacts on the Mire’s resident microbial communities and processes, we sought to link them to their hosts via emerging standard *in silico* host prediction methods, significantly empowered by the recent recovery of 1,529 MAGs from the site (508 from palsa, 588 from bog, and 433 from fen; 16). Tentative bacterial hosts were identified for 17 of the 53 vOTUs (Fig. 3; Table S2): these hosts spanned four genera among three phyla (*Verrucomicrobia: Pedosphaera, Acidobacteria: Acidobacterium* and *Candidatus* Solibacter, and *Deltaproteobacteria: Smithella*). Eight viruses were linked to more than one host, but always within the same species. The four predicted microbial hosts are among the most abundant in the microbial communities, and have notable roles in C cycling (15; 16). Three are acidophilic, obligately aerobic chemoorganoheterotrophs and include the Mire’s dominant polysaccharide-degrading lineage (*Acidobacteria*), and the fourth is an obligate anaerobe shown to be syntrophic with methanogens (*Smithella). Acidobacterium* is a highly abundant, diverse, and ubiquitous soil microbe (85–87), and a member of the most abundant phylum in Stordalen Mire. The relative abundance of this phylum peaked in the bog at 29%, but still had a considerably high relative abundance in the other two habitats (5% in palsa and 3% fen) (16). It is a versatile carbohydrate utilizer, and has recently been identified as the primary degrader of large polysaccharides in the palsa and bog habitats in the Mire, and is also an acetogen (16). Seven vOTUs were inferred to infect *Acidobacterium*, implicating these viruses in directly modulating a key stage of soil organic matter decomposition. The second identified *Acidobacterial* host was in the newly proposed species *Candidatus* Solibacter usitatus, another carbohydrate degrader (88). The third predicted host was *Pedosphaera parvula*, within the phylum *Verrucomicrobia* which is ubiquitous in soil, abundant across our soils (~3% in palsa and ~7% in bog and fen habitats, based on metagenomic relative abundance; 16), utilizes cellulose and sugars (89–93) and in this habitat, this organism could be acetogenic (16). Lastly, vOTU_28 was linked to the *Deltaproteobacteria Smithella* sp. SDB, another acidophilic chemoorganoheterotroph, but an obligate anaerobe, with a known syntrophic relationship with methanogens (94, 95). Collectively, these virus-host linkages provide evidence for the Mire’s viruses to be impacting the C cycle via population control of relevant C-cycling hosts, consistent with previous results in this system (46) and other wetlands (96).

**Figure 3.**
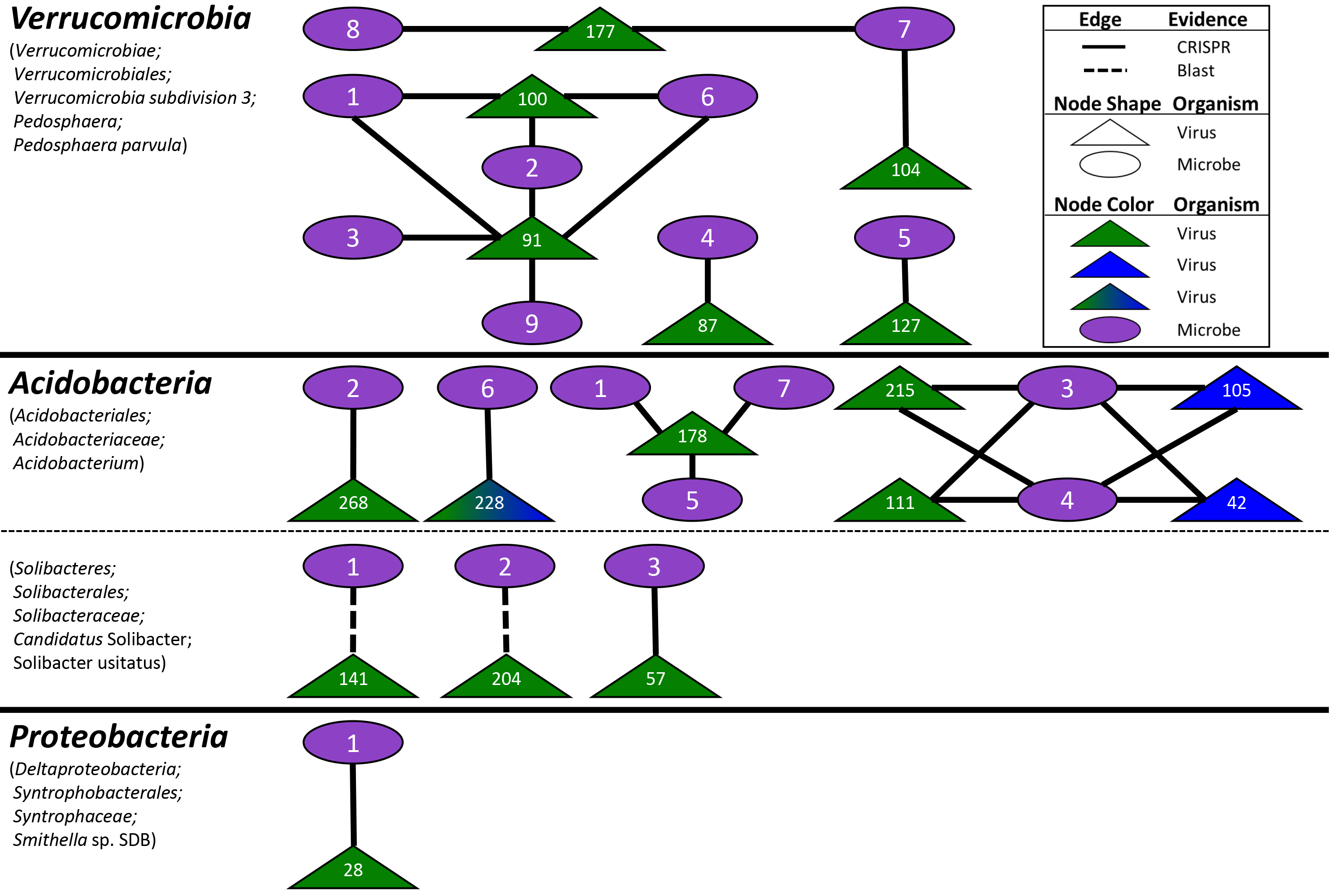
Viral-host linkages between vOTUs and MAGs. Seventeen vOTUs were linked to 4 host lineages by multiple lines of evidence, with 15 linked by CRISPRs (solid line; see Table S2) and 2 by BLAST (dotted line). Node shape denotes organism (oval for microbe and triangle for virus). Viral nodes are color coded by habitat of origin (green for bog and blue for fen).

We next sought to examine viral AMGs for connections to C cycling. To more robustly identify AMGs than the standard protein family-based search approach, we used a custom-built in-house pipeline previously described in Daly et al. (97), and further tailored to identify putative AMGs based on the metabolisms described in the 1,529 MAGs recently reported from these same soils (16). From this, we identified 34 AMGs from 13 vOTUs (Fig. 4; Table S1; Table S3), encompassing C acquisition and processing (three in polysaccharide-binding, one involved in polysaccharide degradation, and 23 in central C metabolism) and sporulation. Glycoside hydrolases that help breakdown complex OM are abundant in resident microbiota (16) and may be especially useful in this high OM environment; notably to our knowledge they have not been found in marine viromes, but have been found in soil (at our site; 46) and rumen (98; Solden et al. *submitted—99*). In addition, central C metabolism genes in viruses may increase nucleotide and energy production during infection, and have been increasingly observed as AMGs (31,32, 33, 34, 35). Finally, two different AMGs were found in regulating endospore formation, *spoVS* and *whiB*, which aid in formation of the septum and coat assembly, respectively, improving spores’ heat resistance (100, 101). A WhiB-like protein has been previously identified in mycobacteriophage TM4 (WhiBTM4), and experimentally shown to not only transcriptionally regulate host septation, but also cause superinfection exclusion (i.e. exclusion of secondary viral infections; 102). While these two sporulation genes have only been found in *Firmicutes* and *Actinobacteria*, the only vOTU to have *whiB* was linked to an acidobacterial host (vOTU_178; Fig. 4). A phylogenic analysis of the *whiB* AMG grouped it with actinobacterial versions and more distantly with another mycobacteriophage (Fig. 4), suggesting either (1) misidentification of host (unlikely, as it was linked to three different acidobacterial hosts, each with zero mismatches of the CRISPR spacer), (2) the virus could infect hosts spanning both phyla (unlikely, as only ~1% of identified virus-host relationships span phyla; 45), or (3) the gene was horizontally transferred into the *Acidobacteria*. Identification of these 34 diverse AMGs (encoded by 25% of the vOTUs) suggests a viral modulation of host metabolisms across these dynamic environments, and supports the findings from bulk metagenome-derived viruses of Emerson et al. (46) at this site. That study’s AMGs spanned the same categories as those reported here, except for *whiB* which was not found, but did not discuss them other than the glycoside hydrolases, one of which was experimentally validated.

**Figure 4.**
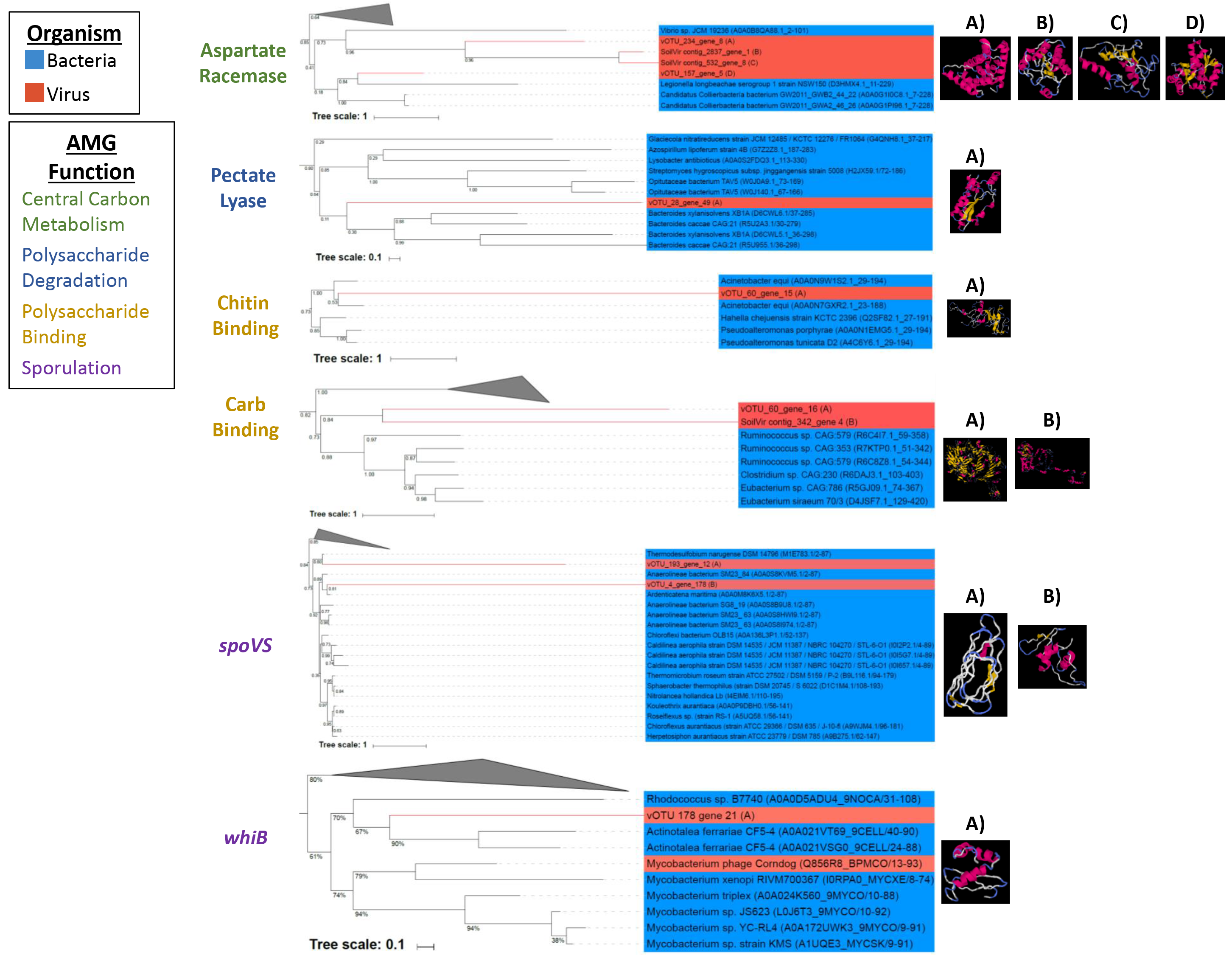
Characterization of select AMGs. FastTree phylogenies were constructed for select AMGs (one from each group and one example for central carbon metabolism), and their structures and those of their nearest neighbors were predicted using iTasser (detailed in Table S3). Tree lineages are shaded blue for bacteria and red for viruses. “vOTU” sequences are from the 53-vOTU virome-derived dataset, while “SoilVir contig” represent homologues from Emerson et al.’s (46) bulk metagenome-derived 1907 vOTUs. AMGs are color coded by function: green for central carbon metabolism, blue for polysaccharide degradation, yellow for polysaccharide binding, and purple for sporulation. The first predicted model for each soil virus is shown and was used for the TM-align comparison. Structures are ordered from left to right based on the appearance of their sequence in the tree from top to bottom.

Thus far, the limited studies of soil viruses have identified few AMGs relative to studies of marine environments. This may be due to under-sampling, or difficulties in identifying AMGs; since AMGs are homologs of host genes, they can be mistaken for microbial contamination (103) and thus are more difficult to discern in bulk-soil metagenomes (whereas marine virology has been dominated by viromes); also, microbial gene function is more poorly understood in soils (104). Alternately, soil viruses could indeed encode fewer AMGs. One could speculate a link between host lifestyle and the usefulness of encoding AMGs; most known AMGs are for photo- and chemo-autotrophs (70, 105, 106), although this may be due to more studies of these metabolisms or phage-host systems. Thus far, soils are described as dominated by heterotrophic bacteria (107–111), and if AMGs were indeed less useful for viruses encoding heterotrophs, that could explain their limited detection in soil viruses. However, a deeper and broader survey of soil viruses will be required to explore this hypothesis.

### Sample storage impacts vOTU recovery

While our previous research demonstrated that differing storage conditions (frozen versus chilled) of these Arctic soils did not yield different viral abundances (by direct counts; 41), the impact of storage method on viral community structure was unknown. Here, we examined that in the palsa and bog habitats for which viromes were successful from both storage conditions. Storage impacted recovered community structure only in the bog habitat, with dramatically broader recovery of vOTUs from the chilled sample (Fig. 5A/B), leading to higher diversity metrics (Fig. S4), and appreciable separation of the recovered chilled-vs-frozen bog vOTU profiles in ordination (Fig. 5C). The greater vOTUs recovery from the chilled sample was likely partly due to higher DNA input and sequencing depth, which was 107-fold more than bog frozen replicate A (BFA) and 350-fold more than bog frozen replicate B (BFB). This led to 1.6- to 9-fold more reads assembling into contigs (compared to viromes BFA and BFB, respectively; Table 1), and 3.5-9-fold more distinct contigs; while one might expect that as the number of reads increased, a portion would assemble into already-established contigs, that was not observed. This higher proportional diversity in the chilled bog virome relative to the two frozen ones could have several potential causes. Freezing might have decreased viral diversity by damaging viral particles, although these viruses regularly undergo freezing (albeit not with the rapidity of liquid nitrogen). Alternatively, there could be a persistent metabolically active microbial community under the chilled conditions with ongoing viral infections, distinct from those in the field community. Finally, there could have been bog-specific induction of temperate viruses under chilled conditions (since this difference was not seen in the palsa samples). The bog habit is very acidic (pH ~4 versus ~6 in palsa and fen; 10, 46), with a dynamic water table, and each of these has been hypothesized or demonstrated to increase selection for temperate viruses (77, 112–116). In addition, of the 19 vOTUs shared between this study and the bulk-soil metagenome study of Emerson et al. (46; which was likely to be enriched for temperate viruses based on its majority sampling of microbial DNA), 13 were unique to the bog, and of those, 10 were only present in the chilled rather than frozen viromes, and the remaining 3 were enriched in the chilled viromes.

**Figure 5.**
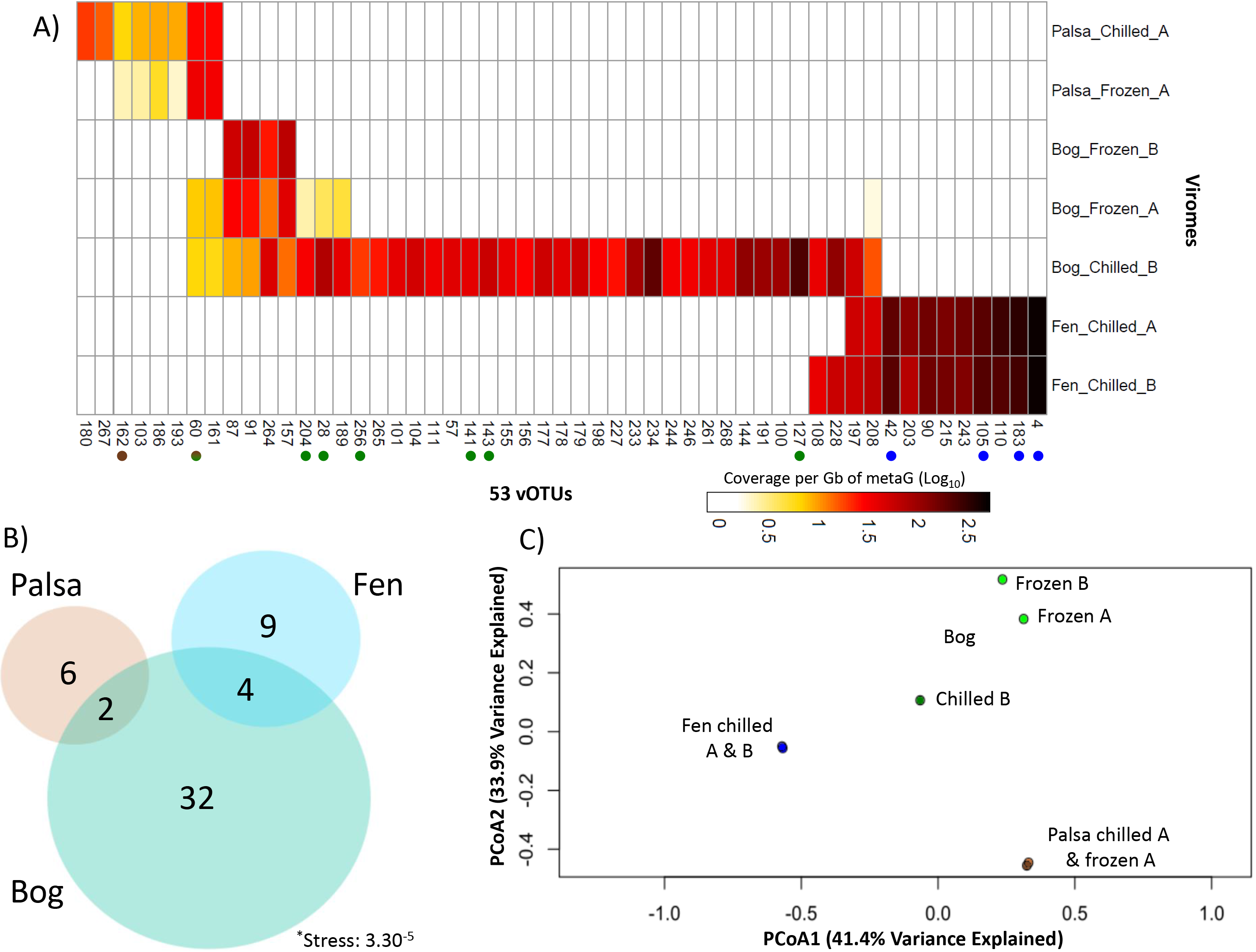
Three views into viral community structure across the thaw gradient. (A) The relative abundance of vOTUs (columns) in the seven viromes (rows). Reads were mapped to this non-redundant set of contigs to estimate their relative abundance (calculated as bp mapped to each read, normalized by contig length and the total bp in each metagenome). Red color gradient indicates log_10_ coverage per Gbp of metagenome. ‘Chilled’ and ‘Frozen’ indicate sample storage at 4°C or flash frozen in liquid nitrogen and stored at ™80°C. “A” and “B” denote technical replicates. Dots after contig names indicate membership in a viral cluster, filled dots denote cluster is novel, and fill color indicates habitat specificity (palsa = brown, bog = green, fen = blue). (B) Euler diagram relating the seven viromes and their 53 vOTUs. (C) Principal coordinate analysis of the viromes by normalized relative abundance of the 53 vOTUs.

Finally, while the chilled bog sample was an outlier to all other viromes (dendrogram, Fig. S5A), a social network analysis of the reads that mapped to the viromes (Fig. S5B & C) indicated that habitat remained the primary driver of recovered communities. Because of this, the diversity analyses were redone with the chilled bog sample taken out (Fig. S2B) instead of subsampling the reads, because this is a smaller dataset (subsampling smaller datasets described further in 117) and the storage effect was observed only for the bog.

### Habitat specificity of the 53 vOTUs along the thaw gradient

We explored the ecology of the recovered vOTUs across the thaw gradients, by fragment recruitment mapping against the (i) viromes, and (ii) bulk-soil metagenomes. Virome mapping revealed that the relative abundance of each habitat’s vOTUs increased along the thaw gradient; relative to the palsa vOTU’s abundances, bog vOTUs were 3-fold more abundant and fen vOTUs were 12-fold more abundant (Fig. 5A). This is consistent with overall increases in viral-like-particles with thaw observed previously at the site via direct counts (41). Only a minority (11%) of the vOTUs occurred in more than one habitat, and none were shared between the palsa and fen (Fig. 5B). Consistent with this, principal coordinates analyses (PCoA; using a Bray-Curtis dissimilarity metric) separated the vOTU-derived community profiles according to habitat type, which also explained ~75% of the variation in the dataset (Fig. 5C). Mapping of the 214 bulk-soil metagenomes from the three habitats (16) revealed that a majority (41; 77%) of the vOTUs were present in the bulk-soil metagenomes (Fig. 6), collectively occurring in 62% (133) of them. Of the 41 vOTUs present, most derived from the bog, and their distribution among the 133 metagenomes reflected this, peaking quite dramatically in the bog (Fig. S4). This strong bog signal in the bulk-soil metagenomes ‒ both in proportion of bog-derived vOTU’s present in the bulk metagenomes, and in abundance of all vOTUs in the bog samples ‒ is consistent with the hypothesized higher abundance of temperate viruses in the bog, suggested by the chilled-versus-frozen storage results above. Overall, vOTU abundances in larger and longer-duration bulk-soil metagenomes indicated less vOTU habitat specificity than in the seven viromes: 10% were unique to one habitat, 22% of vOTUs were present in all habitats, 22% were shared between palsa and bog, 27% between palsa and fen, and 68% between bog and fen (Fig. 6). The difference in observations from vOTU read recruitment of viromes versus bulk-soil metagenomes could be due to many actual and potential differences, arising from their different source material (but from the same sites) and different methodology, including: vOTUs’ actual abundances (they derive from different samples), infection rates, temperate versus lytic states, burst size, and/or virion stability and extractability.

**Figure 6.**
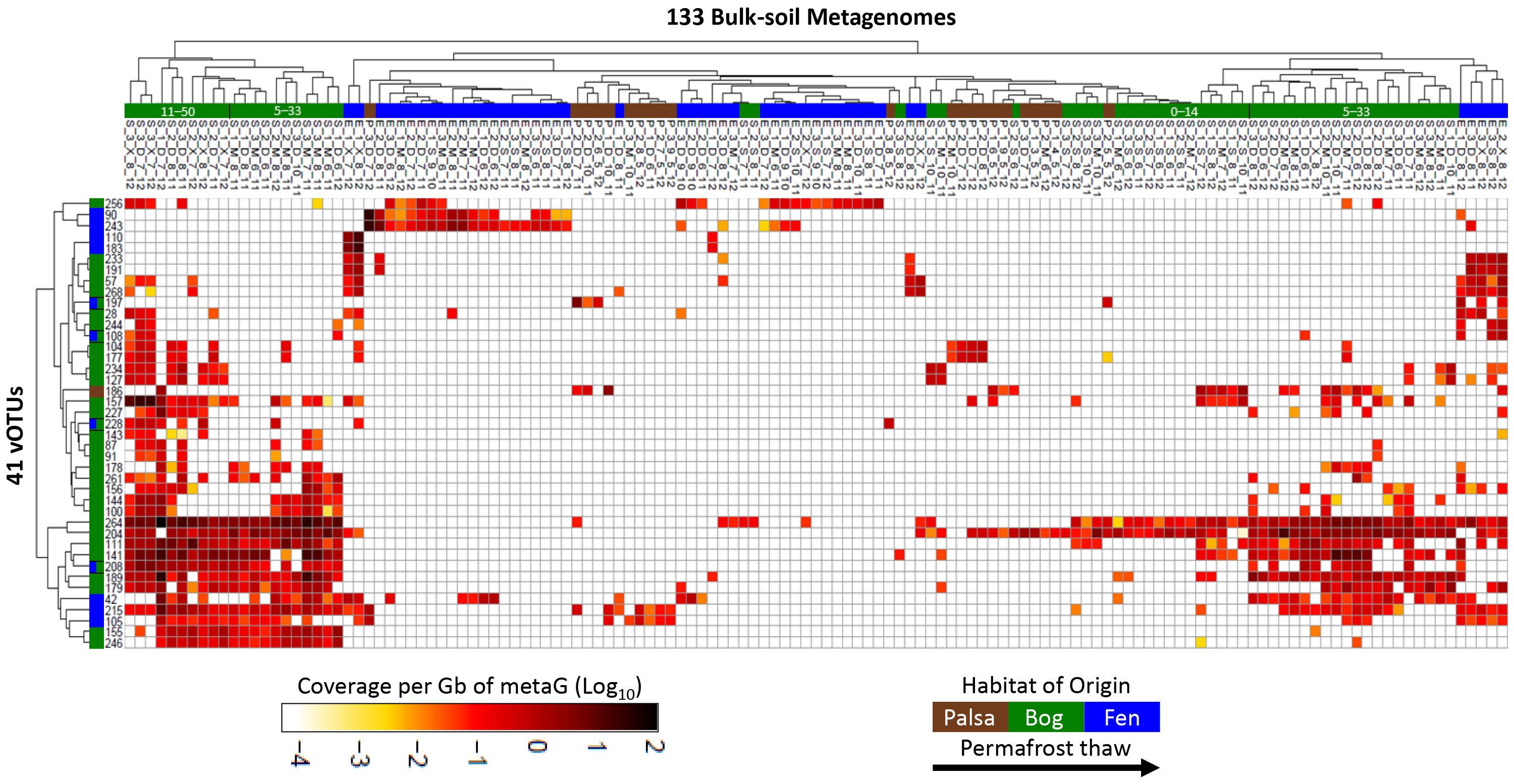
vOTU abundance in 133 bulk-soil metagenomes. The heat map represents abundance of vOTUs (rows) in the bulk-soil metagenomes (columns); metagenome reads were mapped to the non-redundant set of contigs to estimate their relative abundances (calculated as bp mapped to each read, normalized by contig length and the total bp in each metagenome). Red color gradient indicates log10 coverage per Gbp of metagenome. Only the 41 vOTUs present in the metagenomes (out of 53), and the 63 bulk-soil metagenomes (out of 214) that contained matches to the vOTUs, are shown. Metagenome names denote source: habitat of origin (palsa = P, bog = S, fen = E); soil core replicate (1–3); depth (3 cm intervals denoted with respect to geochemical transitions, see 46; generally, S = 1-4 cm, M = 5–14 cm, D = 11–33 cm, and X = 30–50 cm); month collected (5–10 as May-October); year collected (2010, 11, 12). See Emerson et al. (46) and Woodcroft et al. (16) for more metagenome and sample details.

The vOTUs’ habitat preferences observed in both read datasets is consistent with the numerous documented physicochemical and biological shifts along the thaw gradient, and with observations of viral habitat-specificity at other terrestrial sites. Changes in physicochemistry are known to impact viral morphology (reviewed in 37, 118, 119) and replication strategy (36, 37). In addition, at Stordalen Mire (and at other similar sites; 110), microbiota are strongly differentiated by thaw-stage habitat, with some limited overlap among ‘dry’ communities (i.e. those above the water table, the palsa and shallow bog), and among ‘wet’ ones (those below the water table, the deeper bog and fen) (14, 15, 16). These shifting microbial hosts likely impact viral community structure. Expanding from the 53 vOTUs examined here, Emerson et al.’s (46) recent analysis of nearly 2,000 vOTUs recovered from the bulk-soil metagenomes also showed strong habitat specificity among the recovered vOTUs (only 0.1% were shared among all habitats, with <4.5% shared between any two habitats). These findings are also consistent with observations of distinct viral communities from desert, prairie, and rainforests (120), and from grasslands and arctic soils (45). In contrast, an emerging paradigm in the marine field is ‘seascape ecology’ (121), where the majority of taxa are detected across broad geographical areas, as are marine virus (26, 70). This important difference in habitat specificity between soils and oceans may be due to the greater physical structuring of soil habitats.

Although vOTU richness and diversity appeared to increase along the thaw gradient (roughly equivalent in palsa and bog, and ~2-fold higher in fen, omitting the chilled bog sample; Fig. S4), this dataset only captured a small fractioned of the viral diversity (based on the collector’s curve from 46), and therefore the undersampling prevents diversity inferences. Intriguingly, while our virome-derived vOTU richness was lowest in the palsa, Emerson et al.’s (46) much greater sampling recovered the most vOTUs in the palsa, more than double that in the fen (42% vs. 18.9% of total vOTUs). This major difference could potentially be due to the known increase in microbial alpha diversity along the thaw gradient (15, 16), causing increased difficulty of viral genome reconstruction in the bulk-soil metagenomes; specifically, this could be due to poorer assembly of temperate phages within an increasingly diverse microbiota, or of lytic or free viruses due to concomitantly increasing viral diversity (which is consistent with the increased vOTU richness with thaw in our virome dataset). Notably, neither this dataset nor that of Emerson et al. (46) captured ssDNA or RNA viruses, which potentially represent up to half of viral particles (122–124).

### Challenges in characterizing the soil virosphere

The low yield of viral contigs given the relatively large sequencing depth of the viromes reflects several factors that currently challenge soil viromics. First, resuspending viruses from soils is a challenge due to their adsorption to the soil matrix (43). Second, yields of viral DNA are often very low (due to both low input biomass and potentially low extraction efficiency), requiring amplification; this leads to biases (53, 56–61) or poor assembly and few viral contigs (described further in 125). Third, viral contig identification requires a reference database, yet soil viruses are underrepresented in current databases; for example, a majority (85%) of our sequence space was unknown. Fourth, non-viral DNA may co-extract. Lastly, the optimal approach to identifying ecological units within viral sequence space is unclear.

In this study, DNA yields (and sequencing inputs) decreased along the thaw gradient, as did total reads, but counterintuitively viral reads increased (Table 1; the fen had ~5-fold more viral reads than the palsa). This may have been partly due to the shift to a more aquatic-type habitat, for which viruses are better represented in the databases, or to an actual increase in viral DNA (as a portion of total) concomitant with known viral abundance increases (41). A large portion of the assembled reads were non-viral (Table 1), representing either microbial contamination, or gene transfer agents (GTAs), i.e., viral-like capsids that package microbial DNA (reviewed in 126). Since the viral particle purification protocol involved 0.2 μM filter followed by CsCl density gradient-based separation of the viral particles (removing free genomic DNA), contamination by microbial DNA seems unlikely. While ultra-small microbial cells have been found in our soils (46) and other permafrost soils (reviewed in 127, 128), and may have passed through the 0.2 μm filters, they would be expected to be removed in the CsCl gradient since their density is similar to that of larger microbial cells, and not viruses (reviewed in 128–130). Therefore, to identify GTAs we searched our contigs for 16S rRNA genes and for known GTAs. We found six contigs that had 16S rRNA matches to multiple microbes (131), and 94 contigs with matches to known GTAs (126), together accounting for ~25% of the assembled reads. GTAs may thus represent an appreciable and unavoidable ‘contaminant’ in soil viromes, as has been observed in marine systems (reviewed in 126). In this backdrop of potential contaminant DNA and a preponderance of unknown genes in viral sequence space, identifying ecological units in soil viromes is a challenge. We performed a sensitivity analysis on three ways to characterize the ecological units in our dataset: reads characterization, contigs, and as vOTUs (data not shown). While all three methods have validity, there is a higher probability for inclusion of contaminants that can dramatically impact conclusions from the first two approaches. We, therefore, erred on the side of caution and reported our findings in the context of identified vOTUs.

This study’s virome-based approach contrasts (Fig. 7) with that used in Emerson et al. (46), which recovered vOTUs from bulk-soil metagenomes from the same site, but different years, months, depths, and preservation methods. While the viromes derived from separated viral particles, the bulk-soil metagenomes captured viruses within hosts ‒ i.e. those engaged in active infection, and those integrated into hosts ‒ as well as free viruses successfully extracted with the general extraction protocol. This study generated 18 Gb of sequence from 7 viromes, while Emerson et al. (46) analyzed 178 Gb from 190 bulk metagenomes, and neither approach captured the total viral diversity in these soils based on rarefaction (46). The efficiency of vOTU recovery was, unsurprisingly, >2-fold higher using the virome approach (2.93 vOTUs/Gbp of virome, versus 1.30 vOTUs/Gbp of bulk-soil metagenome), suggesting that equivalent virome-focused sequencing effort could yield >4,300 vOTUs (although diversity would likely saturate below that). Of the 19 vOTUs that were shared between the two datasets, the longer, virome-derived sequences defined them. These findings suggest that viromes (which greatly enrich for viral particles) and bulk-soil metagenomes (which are less methodologically intensive, and provide simultaneous information on both viruses and microbes) can offer complementary views of viral communities in soils and the optimal method will depend on the goal of the study.

**Figure 7.**
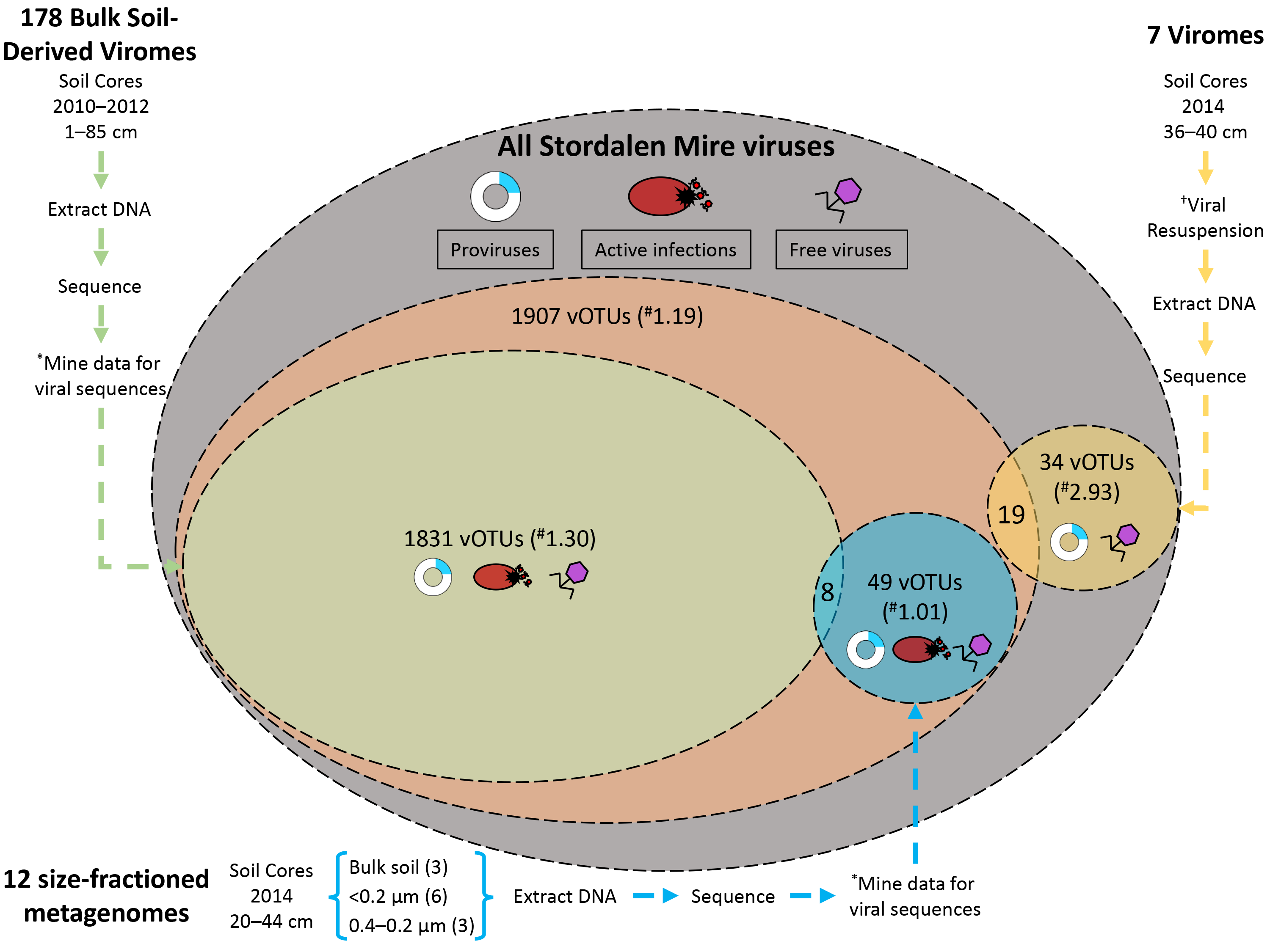
Contrasting Stordalen Mire viruses derived from viromes and bulk-soil metagenomes. Currently, two datasets exist describing Stordalen Mire (SM) Archaeal and Bacterial viruses. Emerson et al. (46) characterized the viral signal in bulk-soil metagenomes (described in 16), while here we characterize viruses from viromes, derived from separated viral particles. There are three possible stages of the viral life cycle at which to capture viruses: proviruses (those integrated into a host genome; blue), active infections (viruses undergoing lytic infection; red), and free viruses (viruses not currently infecting a host; purple). The largest oval represents all the theoretical SM viruses (gray). The next largest oval represents the vOTUs reported in Emerson et al. (46; orange). Within that oval are the vOTUs derived from bulk-soil metagenomes (green), and from size-fractioned bulk-soil metagenomes also used in that study (blue). The final oval represents the vOTUs identified in this study (yellow circle). Nineteen vOTUs are shared between the two datasets (57, 90, 110, 111, 141, 144, 155, 157, 179, 183, 186, 189, 197, 204, 208, 243, 246, 261, and 264). Also shown are the methods that produced each dataset. * denotes the viral signal was mined from bulk-soil metagenomes. † denotes that viruses were resuspended from the soils using a previously optimized protocol (41). # denotes the vOTU yield normalized per Gbp of metagenome. The active viruses or proviruses detected in the size-fractioned bulk-soil metagenomes are only those that infect microbial hosts that could pass through the reduced pore size filters (more sample information in 46).

Over the last 2 decades, viruses have been revealed to be ubiquitous, abundant, and diverse in many habitats, but their role in soils has been underexplored. The observations made here from virome-derived viruses in a model permafrost-thaw ecosystem show these vOTUs are primarily novel, change with permafrost thaw, and infect hosts highly relevant to C cycling. The next important step is to more comprehensively characterize these viral communities (from more diverse samples, and including ssDNA and RNA viruses), and begin quantifying their direct and indirect impacts on C cycling in this changing landscape. This should encompass the complementary information present in virome, bulk metagenomes, and viral signal from MAGs, analyzed in the context of the abundant metadata available. With increasing characterization of soil viruses, their mechanistic interactions with hosts, and quantification of their biogeochemical impacts, soil viral ecology may significantly advance our understanding of terrestrial ecosystem biogeochemical cycling, as has marine viral ecology in the oceans.

## Methods and Materials

### Sample collection

Samples were collected from July 16–19, 2014 from peatland cores in the Stordalen Mire field site near Abisko, Sweden (Fig. 1; more site information in 7, 10, 12). The soils derived from palsa (one stored chilled and the other stored frozen), bog (one stored chilled and two stored frozen), and fen (both stored chilled) habitats along the Stordalen Mire permafrost thaw gradient. These three sub-habitats are common to northern wetlands, and together cover ~98% of Stordalen Mire’s non-lake surface (8). The sampled palsa, bog, and fen are directly adjacent, such that all cores were collected within a 120 m total radius. For this work, the cores were subsampled at 36–40 cm, and material from each was divided into two sets. Set 1 was chilled and stored at 4°C, and set 2 was flash-frozen in liquid nitrogen and stored at −80°C as described in Trubl et al. (41). Both sets were processed using a viral resuspension method optimized for these soils (41). For CsCl density gradient purification of the particles, CsCl density layers of rho 1.2, 1.4, 1.5, and 1.65 were used to establish the gradient; we included a 1.2 g/cm^3^ CsCl layer to try to remove any small microbial cells that might have come through the 0.2um filter (for microbial cell densities see 132, 133; for viral particle densities see 50). We then collected the 1.4-1.52 g/cm^3^ range from the gradient for DNA extraction, to target the dsDNA range (per 50). The viral DNA was extracted using Wizard columns (Promega, Madison, WI, products A7181 and A7211), and cleaned up with AMPure beads (Beckman Coulter, Brea, CA, product A63881). DNA libraries were prepared using Nextera XT DNA Library Preparation Kit (Illumina, San Diego, CA, product FC-131-1024) and sequenced using an Illumina MiSeq (V3 600 cycle, 6 samples/run, 150 bp paired end) at the University of Arizona Genetics Core facility (UAGC). Seventeen viral contigs were previously described in Emerson et al. (46) (Fig. 7).

The 214 bulk-soil metagenomes and associated recovered MAGs used here for analyses were described in Woodcroft et al. (16), and derive from the same sampling sites from 2010-2012, and 1–85 cm depths. They were extracted using a modification of the PowerSoil kit (Qiagen, Hilden, Germany) and sequenced via TruSeq Nano (Illumina) library preparation or for low concentration DNA samples, libraries were created using the Nextera XT DNA Sample Preparation Kit (Illumina), as described in Woodcroft et al (16).

### vOTU recovery

Eight viromes were prepped and seven samples were successfully sequenced (2 palsa: one chilled and one frozen; 3 bog: one chilled and two frozen; and 2 fen: both chilled). The sequences were quality-controlled using Trimmomatic (134; adaptors were removed, reads were trimmed as soon as the per-base quality dropped below 20 on average on 4 nt sliding windows, and reads shorter than 50 bp were discarded), then assembled separately with IDBA-UD (135), and contigs were processed with VirSorter to distinguish viral from microbial contigs (virome decontamination mode; 66). The same contigs were also compared by BLAST to a pool of putative laboratory contaminants (i.e. phages cultivated in the lab: *Enterobacteria* phage PhiX17, Alpha3, M13, *Cellulophaga baltica* phages, and *Pseudoalteromonas* phages). All contigs matching these genomes at more than 95% average nucleotide identity (ANI) were removed. VirSorted contigs were manually inspected by observing the key features of the viral contigs that VirSorter evaluates (e.g. the presence of a viral hallmark gene places the contigs in VirSorter categories 1 or 2, but further inspection is needed to confirm it is a genuine viral contig and not a GTA or plasmid). To identify GTAs we searched through all of our contigs assembled by IDBA-UD for (1) taxa related to the 5 types of GTAs (keyword searches were: *Rhodobacterales*, *Desulfovibrio*, *Brachyspira*, *Methanococcus*, and *Bartonella*) and (2) microbial DNA the SILVA ribosomal RNA database (release 128; 131), with all the assembled contigs with ≥95% ANI. The percent of reads that mapped to these contigs was calculated as previously described.

After having verified that the VirSorted contigs were genuine viruses, quality controlled reads from the seven viromes were pooled and assembled together with IDBA-UD to generate a non-redundant set of contigs. Resulting contigs were re-screened as described above, removing all identifiable contamination. The contigs then underwent further quality checks by (i) removing all contigs <10 kb and (ii) only using contigs from VirSorter categories 1 and 2.

To detect putative archaeal viruses, the VirSorter output was used as an input for MArVD (with default settings; 136). The output putative archaeal virus sequences were then filtered to include only those contigs ≥10 kb in size resulting in the set of putative archaeal vOTUs described here.

Viral genes were annotated using a pipeline described in Daly et al. (97). Briefly, for each contig, ORFs were freshly predicted using MetaProdigal (137) and sequences were compared to KEGG (138), UniRef and InterproScan (139) using USEARCH (140), with single and reverse best-hit matches greater than a 60 bitscore. AMGs were identified by manual inspection of the protein annotations guided by known resident microbial metabolic functions (identified in 16). To determine confidence in functional assignment, representatives for each AMGs underwent phylogenetic analyses. First each sequence was BLASTed and the top 100 hits were investigated to identify main taxa groups. An alignment with the hits and the matching viral sequence (MUSCLE with default parameters; 141) was done with manual curation to refine the alignment (e.g. regions of very low conservation from the beginning or end were removed). FastTree (default parameters with 1000 bootstraps; 142) was used to make the phylogeny and iTol (143) was used to visualize and edit the tree (any distance sequences were removed). To see if this AMG was wide-spread across the putative soil viruses, a BLASTp (default settings) of each AMG against all putative viral proteins from our viromes was done. The sequences from identified homologs (based on a bitscore >70 and an e value of 10^−4^) were used with the AMG of interest to construct a new phylogenic tree (same methods as before). Finally, structures were predicted using i-TASSER (144) for our AMGs of interest and their neighbors. To assess correct structural predictions, AMGs of interest and their neighbors’ structures were compared with TM-align (TM-score normalized by length of the reference protein; 145).

### Gene-sharing network construction, analysis, and clustering of viral genomes (fragments)

We built a gene-sharing network, where the viral genomes and contigs are represented by nodes and significant similarities as edges (71, 72). We downloaded 198,556 protein sequences representing the genomes of 1,999 bacterial and archaeal viruses from NCBI RefSeq (v 75; 146). Including protein sequences from the 53 Stordalen Mire viral contigs, a total of 199,613 protein sequences were subjected to all-to-all BLASTp searches, with an e-value threshold of 10^−4^, and defined as protein clusters (PCs) in the same manner as previously described (67). Based on the number of PCs shared between the genomes and/or genome fragments, a similarity score was calculated using vConTACT (71, 72). The resulting network was visualized with Cytoscape (version 3.1.1; http://cytoscape.org/), using an edge-weighted spring embedded model, which places the genomes or fragments sharing more PCs closer to each other. 398 RefSeq viruses not showing significant similarity to viral contigs were excluded for clarity. The resulting network was composed of 1,722 viral genomes including 53 contigs and 58,201 edges. To gain detailed insights into the genetic connections, the network was decomposed into a series of coherent groups of nodes (aka VCs; 69, 71, 72), with an optimal inflation factor of 1.6. Thus, the discontinuous network structure of individual components, together with the isolated contigs, indicates their distinct gene pools (68). To assign contigs into VCs, PCs needed to include ≥2 genomes and/or genome fragments, then Markov clustering (MCL) algorithm was used and the optimal inflation factor was calculated by exploring values ranging from 1.0 to 5 by steps of 0.2. The taxonomic affiliation was taken from the NCBI taxonomy (http://www.ncbi.nlm.nih.gov/taxonomy).

### vOTU ecology

Virome reads were mapped back to the non-redundant set of contigs to estimate their coverage, calculated as number of bp mapped to each read normalized by the length of the contig, and by the total number of bp sequenced in the metagenome in order to be comparable between samples (Bowtie 2, threshold of 90% average nucleotide identity on the read mapping, and 75% of contig covered to be considered as detected; 54, 147). The heat map of the vOTU’s relative abundances across the seven viromes, as inferred by read mapping, was constructed in R (CRAN 1.0.8 package pheatmap).

The 214 bulk-soil metagenomes and 1,529 associated recovered MAGs used here for analyses were described in Woodcroft et al. (16). The paired MAG reads were mapped to the viral contigs with Bowtie2 (as described above for the virome reads). The heat map of the vOTU’s relative abundances across the 214 bulk-soil metagenomes, as inferred by read mapping, was constructed in R (CRAN 1.0.8 package pheatmap); only microbial metagenomes with a viral signal were shown.

### Viral-host methodologies

We used two different approaches to predict putative hosts for the vOTUs: one relying on CRISPR spacer matches (45, 97, 148) and one on direct sequence similarity between virus and host genomes (149). For CRISPR linkages, Crass (v0.3.6, default parameters), a program that searches through raw metagenomic reads for CRISPRs was used (further information in Table S2; 150). For BLAST, the vOTU nucleotide sequences were compared to the MAGs (16) as described in Emerson et al. (46). Any viral sequences with a bit score of 50, E-value threshold of 10^−3^, and ≥70% average nucleotide identity across ≥2500 bp were considered for host prediction (described in 151).

### Phylogenetic analyses to resolve taxonomy

Two phylogenies were constructed. The first had the alignment of the protein sequences that are common to all *Felixounavirinae* and *Vequintavirinae* as well as vOTU_4 and the second had an alignment of select sequences from PC_03881, including vOTU_165. These alignments were generated using the ClustalW implementation in MEGA5 (version 5.2.1; http://www.megasoftware.net/). We excluded non-informative positions with the BMGE software package (152). The alignments were then concatenated into a FASTA file and the maximum likelihood tree was built with MEGA5 using JTT (jones-Taylor-Thornton) model for each tree. A bootstrap analysis with 1,000 replications was conducted with uniform rates and a partial depletion of gaps for a 95% site coverage cutoff score.

### Accession numbers

All data (sequences, site information, supplemental tables and files) are available as a data bundle at the IsoGenie project database under data downloads at https://isogenie.osu.edu/. Additionally, viromes were deposited under BioProject ID PRJNA445426 and SRA SUB3893166, with the following BioSample accession numbers: SAMN08784142 for Palsa chilled replicate A, SAMN08784143 for Palsa frozen replicate A, SAMN08784152 for Bog frozen replicate A, SAMN08784154 for Bog frozen replicate B, SAMN08784153 for Bog chilled replicate B, SAMN08784163 for Fen chilled replicate A, and SAMN08784165 for Fen chilled replicate B.

## Acknowledgments

We thank Bonnie Poulos and Christine Schirmer for their assistance on different stages of this project. We also thank SWES-MEL, TMPL, and The University of Arizona Genetics Core facility, MAVERIC lab at the Ohio State University, the Abisko Naturvetenskapliga Station, and the Joint Genome Institute for support. We thank Moira Hough, Robert Jones, and Rachel Wilson for sample collection assistance. Bioinformatics were supported by The Ohio Supercomputer Center and by the National Science Foundation under Award Numbers DBI-0735191 and DBI-1265383; URL: www.cyverse.org. This study was funded by the Genomic Science Program of the United States Department of Energy Office of Biological and Environmental Research, (grants DE-SC0004632, DE-SC0010580, and DE-SC0016440), and by a Gordon and Betty Moore Foundation Investigator Award (GBMF#3790 to MBS). We thank Dr. Michael Palace for generating and allowing us to use the unmanned aerial vehicle (UAV) image in Fig. S1.

## Table legends

**Table 1. Soil viromes read information.** The seven viromes are provided, along with their DNA quantity, total number of reads, total number of assembled reads, the number of reads that mapped to soil viral contigs, the number of reads that mapped to the 53 vOTUs, and the average adjusted coverage. Adjusted coverage was calculated by mapping reads back to this non-redundant set of contigs to estimate their relative abundance, calculated as number of bp mapped to each read normalized by the length of the contig and the total number of bp sequenced in the metagenome. For a read to be mapped it had to have ≥90% average nucleotide identity between the read and the contig, and then for a contig to be considered as detected reads had to cover ≥75% of the contig.

**Table 2. Soil viruses’ bioinformatics information.** All 393 putative soil viruses are listed (378 after VirSorter/MArVD and manual inspection). For the vOTUs, the virome(s) in which it originated from, its genomic information, and its coverage is provided. For the other putative soil viral contigs, the origin virome(s) is provided, and contig length are provided. Additionally, the three mobile genetic elements and ten viral contigs with no coverage are reported with their virome(s) of origin (if applicable) and contig length. No contigs were chimeric (i.e. constructed with reads coming from multiple viromes). A † denotes the contig did not meet our threshold for read mapping (i.e. reads recruited to contigs only if they had 90% ANI and then if ≥ 70% of the contig was covered) and therefore could not be counted as detected.

## Supplementary Table legends

**Table S1. Virally-encoded auxiliary metabolic genes and other genes of interest.** Genes were annotated and AMGs identified by running assembled contigs through a pipeline developed by the Wrighton lab at The Ohio State University previously described in Daly et al. (23). The habitat that the vOTU was derived from is listed. Predicted genes that are AMGs or integrase-related are bolded and unannotated genes are no present. Additionally, the PhoH-like protein is bolded due to its highly debated function as a phosphate starvation gene (reviewed in 24).

**Table S2. Viral-host linkages supporting information.** NCBI BLAST linkages were determined based on queries and CRISPR information was provided using Crass software. Host genomes IDs were assigned from the Joint Genome Institute’s Integrated Microbial Genomes Database. Microbial bins were pulled from Woodcroft et al. (*1*).

**Table S3. Structural comparison between select AMGs and phylogenetic neighbors.** Predicted structures for AMGs and neighbors were determined and a comparison of the first model of their predicted structure was performed using TM-align. Structure similarity between two proteins is rated on a scale of 0.0–1.0, with TM-scores < 0.30 suggest random structural similarity and scores = 0.5 suggest similar folds and scores near 1 suggest a perfect match between two structures.

**Table S4. Codon usage frequency.** The codon usage frequency was determined for the 53 vOTUs and the linked microbial bins.

## Supplementary Figure legends

**Figure S1. Phylogenetic analysis of vOTU 4.** Phylogenetic relationships between vOTU_4 and its related viruses. A maximum-likelihood tree was constructed upon a concatenation of two structural proteins (major capsid protein and baseplate protein) that are common to the *Felixounvirinae* and *Vequintavirinae* viruses. The numbers at the branch represent the bootstrapping probabilities from 1000 replicates. Edges with bootstrap values above 75% are represented. The scale bar indicates the number of substitution per site.

**Figure S2. Viral biodiversity increases with permafrost thaw.** Richness, Shannon’s Diversity index and Pielou’s evenness index were calculated for each virome and the viromes were plotted by habitat. Chilled samples are denoted with a lighter color and frozen samples denoted with a darker color. (A) The diversity indices for all seven viromes. (B) The diversity indices of six viromes (bog chilled B was removed).

**Figure S3. Visualizing relationships among soil viral communities.** The y-axis is a measure of Bray-Curtis dissimilarity, with an average dissimilarity used for viromes (i.e. dissimilarities are averaged at each step between viromes for the agglomerative method). Bootstraps n=1000; two types of *p*-values: Approximately Unbiased (AU) *p*-value in blue and Bootstrap Probability (BP) value in purple. AU *p*-value, which is computed by multiscale bootstrap resampling, is a better approximation to unbiased p-value than BP value computed by normal bootstrap resampling. Social networks of: (A) the 53 vOTU sequences and (B) all the reads mapped to the 53 vOTUs from the seven viromes with clusters circled in black. Dots in the social networks represent statistical samples taken from the marginal posterior distributions (Bayesian Method).

**Figure S4. Identified Viral Signal in the MAGs.** (A) The stacked bar chart shows the percent of viral signal occurrences from the 53 vOTUs collected in 2014 in the 133 bulk-soil metagenomes that had a signal collected from 2010-2012. In 2010, only fen samples were collected for microbial metagenomes. Viral signal occurrences were normalized by the number of viromes constructed for each habitat and the number of metagenomes for each habitat. The total number of occurrences for each year is italicized. (B) The number of occurrences (presented as a percentage) of a ‘viral signal’ in a bulk-soil metagenome partitioned by the origin of the bulk-soil metagenome and the vOTU.

**Figure S5. Codon usage frequency for the linked viruses and their microbial hosts.** Principal Coordinates Analysis of the codon usage frequency of microbial hosts and their linked viruses, using the Bray-Curtis dissimilarity metric. Microbial hosts are denoted by circles and colored by phylum (A) or genus/species (B). The associated viruses have a matching color to its host and are denoted with a square. (C) The average dissimilarity metric between the viral contigs linked to potential microbial hosts is plotted against each viruses’ contig length (x10^3^). Average dissimilarity distance was used with viral contigs with multiple hosts.

